# FBXO38 does not control PD-1 stability

**DOI:** 10.1101/2023.03.19.531556

**Authors:** Nikol Dibus, Eva Salyova, Karolina Kolarova, Michele Pagano, Ondrej Stepanek, Lukas Cermak

**Affiliations:** Laboratory of Cancer Biology, Institute of Molecular Genetics of the Czech Academy of Sciences, Czech Republic; Laboratory of Adaptive Immunity, Institute of Molecular Genetics of the Czech Academy of Sciences, Prague, Czech Republic; Department of Biochemistry and Molecular Pharmacology, New York University Grossman School of Medicine, New York, NY 10016, USA; Laura and Isaac Perlmutter NYU Cancer Center, New York University Grossman School of Medicine, New York, NY 10016, USA; Howard Hughes Medical Institute, New York University Grossman School of Medicine, New York, NY 10016, USA

**Keywords:** PD-1, FBXO38, Cullin, immune checkpoint, protein degradation

## Abstract

SKP1-CUL1-F-box protein (SCF) ubiquitin ligases are versatile protein complexes that mediate the ubiquitination of substrates, which are recognized by their F-box-domain-containing subunits^1^. One of these substrate receptors is FBXO38. Its gene has been found to be mutated in several families with early-onset distal hereditary motor neuronopathy^2^. SCF^FBXO38^ ubiquitin ligase controls the stability of ZXDB, a nuclear factor associated with the centromeric chromatin protein CENP-B^3^. Moreover, the loss of FBXO38 results in growth retardation and defect in spermatogenesis characterized by deregulation of the Sertoli cell transcription program and centromere integrity^4^. A report by Meng et al. proposed that SCF^FBXO38^ regulates the protein levels of the PD-1 inhibitory receptor (also known as CD279, PDCD1) in T cells^5^. Here, we have re-addressed the conclusions by Meng et al. using *Fbxo38*^*KO/KO*^ mice and cell systems. We have found no evidence indicating that FBXO38 controls the abundance and stability of PD-1.

## Results

We initially focused on two main aspects presented in the report by Meng et al.: (i) the physical interaction between FBXO38 and PD-1 and (ii) the stability of PD-1 protein in the absence of FBXO38.

In most of their biochemical experiments, Meng et al. overexpressed both FBXO38 and PD-1 in HEK293FT cells. We noticed that PD-1 overexpression in HEK293FT cells led to a significant accumulation of PD-1 protein in the cytoplasm (Extended Data Fig. 1a; white arrowheads). To avoid artifacts due to overexpression-mediated aggregation, we used a Sleeping beauty transposon-based system expressing human PD-1 at near physiologic levels upon doxycycline induction (HEK293FT-PD-1). Moreover, instead of using tagged PD-1, we optimized the use of a commercially available antibody to detect PD-1 protein in western blots and paraformaldehyde-fixed cells (Extended Data Fig. 1a,b). This allowed us to determine whether FBXO38 and PD-1 are located in the same cellular compartments. As with endogenous PD-1, doxycycline induction led to PD-1 expression predominantly at the plasma membrane (Fig. 1a)^6^. Our previous studies demonstrated that FBXO38 is localized in the nucleus both in cultured cells and *in vivo*^3,4^. This is a consequence of its strong nuclear localization signals. To test whether PD-1 expression affects FBXO38 localization or whether PD-1 could be stabilized in the nucleus, we detected both FBXO38 and PD-1 by immunofluorescence upon inhibition of the proteasome (with MG-132) or inhibition of neddylation (using MLN4924, which prevents SCF-mediated degradation)^7^. In contrast to plasma membrane-associated PD-1, endogenous FBXO38 was strictly nuclear. Incubation with MG-132 or MLN4924 did not lead to co-localization of FBXO38 and PD-1 in any subcellular compartment (Fig. 1a and Extended Data Fig. 1c).

**Fig. 1.**
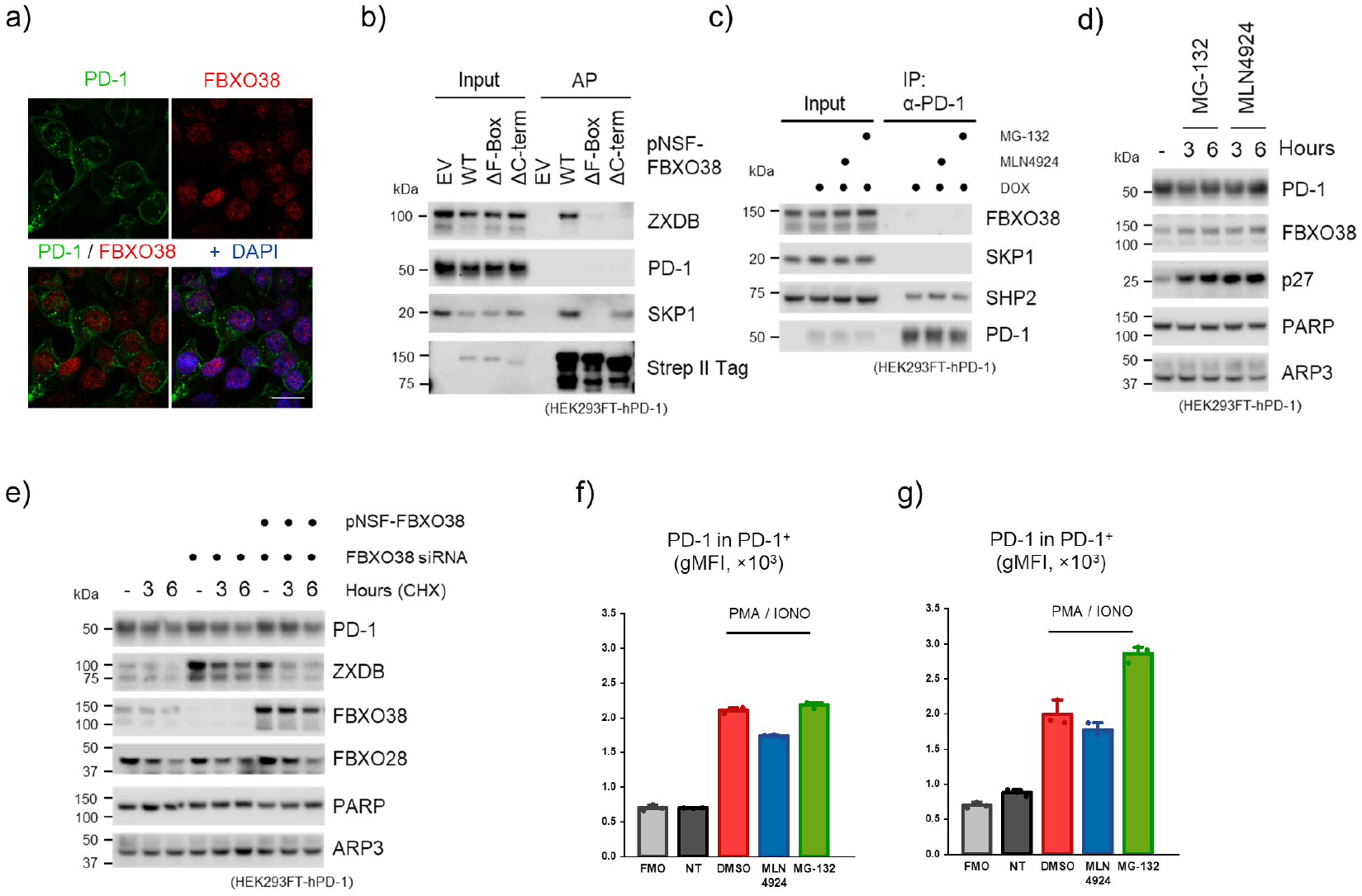
FBXO38 does not control PD-1 in the cell system. (a) FBXO38 does not colocalize with PD-1. HEK293FT cells with the inducible expression of human PD-1 (HEK293FT-hPD-1) were incubated with doxycycline for 48 h and subsequently fixed, permeabilized, and incubated with anti-FBXO38 and PD-1 antibodies. DNA was stained with DAPI. Scale bar, 20 µm. (b) HEK293FT-hPD-1 cells were transfected with StrepII-FLAG-tagged wild-type FBXO38 (pNSF-FBXO38 WT), FBXO38 lacking the F-box motif (ΔF-Box) or C-terminus (ΔC-term). The cells were treated with MLN4924 inhibitor 6 h prior to collecting, lysed, and subjected to affinity purification (AP) using Strep-Tactin resin and immunoblotted as indicated. SKP1 or ZXDB staining was used as the positive control. Inputs represent 1 % of the whole-cell lysates subjected to AP. (c) HEK293FT-hPD-1 cells were treated with doxycycline, MG-132, and MLN4924 where stated. Whole-cell lysates were subjected to immunoprecipitation (IP) and immunoblotted as indicated. SHP2 staining was used as a positive control. Inputs represent 1 % of the whole-cell lysate subjected to IP. (d) HEK293FT-hPD-1 cells were treated with MLN4924 and MG-132 for 3 or 6 h, lysed, and immunoblotted as indicated. p27 staining was used as a positive control, and PARP or ARP3 as the loading control. (e) HEK293FT-hPD-1 cells were transfected with non-targeting siRNA or a mixture of three different siRNAs targeting FBXO38 along with an empty pcDNA3.1 vector or SF-FBXO38 WT vector. Cells were then treated with cycloheximide (CHX) for the indicated time. Whole- cell lysates were immunoblotted as indicated. ZXDB or FBXO28 staining was used as the positive control, and PARP or ARP3 as loading controls. (f,g) Jurkat (f) or HPB-ALL (g) T-cell leukemia cell lines were incubated for 72 h with phorbol myristyl acetate (PMA) and ionomycin (IONO) followed by 6 h treatments with MG-132 or MLN4924 (*n=* 3). Surface PD-1 expression was analyzed by flow cytometry. Bars represent the geometric mean of PD-1 fluorescence intensities in the PD-1^+^ population (gMFI + SD).

Next, we tested the interaction between FBXO38 and PD-1. We pretreated the cells with (MLN4924) to avoid the degradation of PD-1, which could be potentially caused by FBXO38 overexpression. Consistent with our previous observations^3,4^, FBXO38 interacted with its substrate ZXDB, and deletion of either its C-terminus or its F-box motif disrupted this interaction (Fig. 1b). However, we did not detect any interaction between PD-1 and FBXO38. To verify that inducible PD-1 correctly folds and associates with its common interactors^8^, we immunopurified it from HEK293FT-PD-1 cells. Immunoprecipitated PD-1 interacted with its canonical partner SHP2, but not with FBXO38 or SKP1 (Fig. 1c).

Moreover, the inhibition of the proteasome- or SCF-dependent degradation did not affect PD-1 protein levels, while increasing the protein level of p27, an established SCF^FBXL1^ substrate (Fig. 1d)^9^. Meng et al. overexpressed FBXO38 in HEK293FT cells to enhance PD-1 degradation. However, we have previously shown that FBXO38 overexpression does not result in forced degradation of its substrate^3^. Accordingly, we observed only a modest decrease in ZXDB protein level and no changes in PD-1. Inhibition of SCF-dependent degradation in the presence of FBXO38 led to an increase in ZXDB and p27 levels, with no effects on PD-1 protein levels (Extended Data Fig. 1d).

We also evaluated the half-life of PD-1. To this end, we monitored PD-1 protein levels before and after inhibiting protein translation by cycloheximide in control and FBXO38- depleted cells. We observed a significant increase in ZXDB protein levels and stability in FBXO38 knockdown cells, but no changes in PD-1 (Fig. 1e). FBXO38 overexpression restored ZXDB degradation, but exhibited no effect on PD-1 protein levels.

PD-1 is a T-cell-specific gene, so, to rule out the possibility that our observations could be limited to epithelial cell lines such as HEK293FT, we followed Meng et al.’s approach and assessed PD-1 levels in T-cell leukemia cell lines and primary mouse T-cells. First, we induced the expression of endogenous PD-1 in Jurkat and HPB-ALL cell lines using ionomycin and phorbol myristyl acetate (PMA) in the presence and in the absence of proteasome and neddylation inhibitors. Flow cytometry analysis showed that upon activation, the majority of the cells exhibited strong surface PD-1 expression (Extended Data Fig. 1e,f). In contrast to Jurkat cells, where proteasome inhibition had no significant effect, in the HPB-ALL cell line, treatment with MG-132 increased the percentage of PD-1 positive cells and the level of PD-1 surface expression (Figure 1f,g and Extended Data Fig. 1f,g). Importantly, inhibition of SCF-dependent degradation in both cell lines led to a decrease in PD-1 surface expression and the percentage of PD-1 positive cells (Fig. 1f,g and Extended Data Fig. 1f,g), rather than their increase, as it would be expected based on the claims by Meng et al.

To evaluate the effect of FBXO38 deficiency on PD-1 expression, cell-surface levels of PD-1 in T cells from WT and *Fbxo38*^*KO/KO*^ mice were analyzed by flow cytometry at steady state and upon activation. At steady state, PD-1 was expressed almost exclusively in regulatory CD4^+^ CD25^+^ T cells at similar levels in both genotypes (Fig. 2a and Extended Data Fig. 2a). *Ex vivo* activation with anti-CD3/CD28 beads in the presence of IL-2 induced the upregulation of PD-1 surface levels in CD4^+^ and CD8^+^ T cells (Fig. 2b). However, *Fbxo38*^*KO/KO*^ T cells did not exhibit higher PD-1 surface levels than their WT counterparts, as it would have been expected if PD-1 was a substrate for SCF^FBXO38^. If anything, the PD-1 surface levels were rather slightly lower in *Fbxo38*^*KO/KO*^ than in WT T cells. In contrast, the SCF^FBXO38^ substrate ZXDB was upregulated in *ex vivo* activated *Fbxo38*^*KO/KO*^ T cells (Extended Data Fig. 2b).

**Fig. 2.**
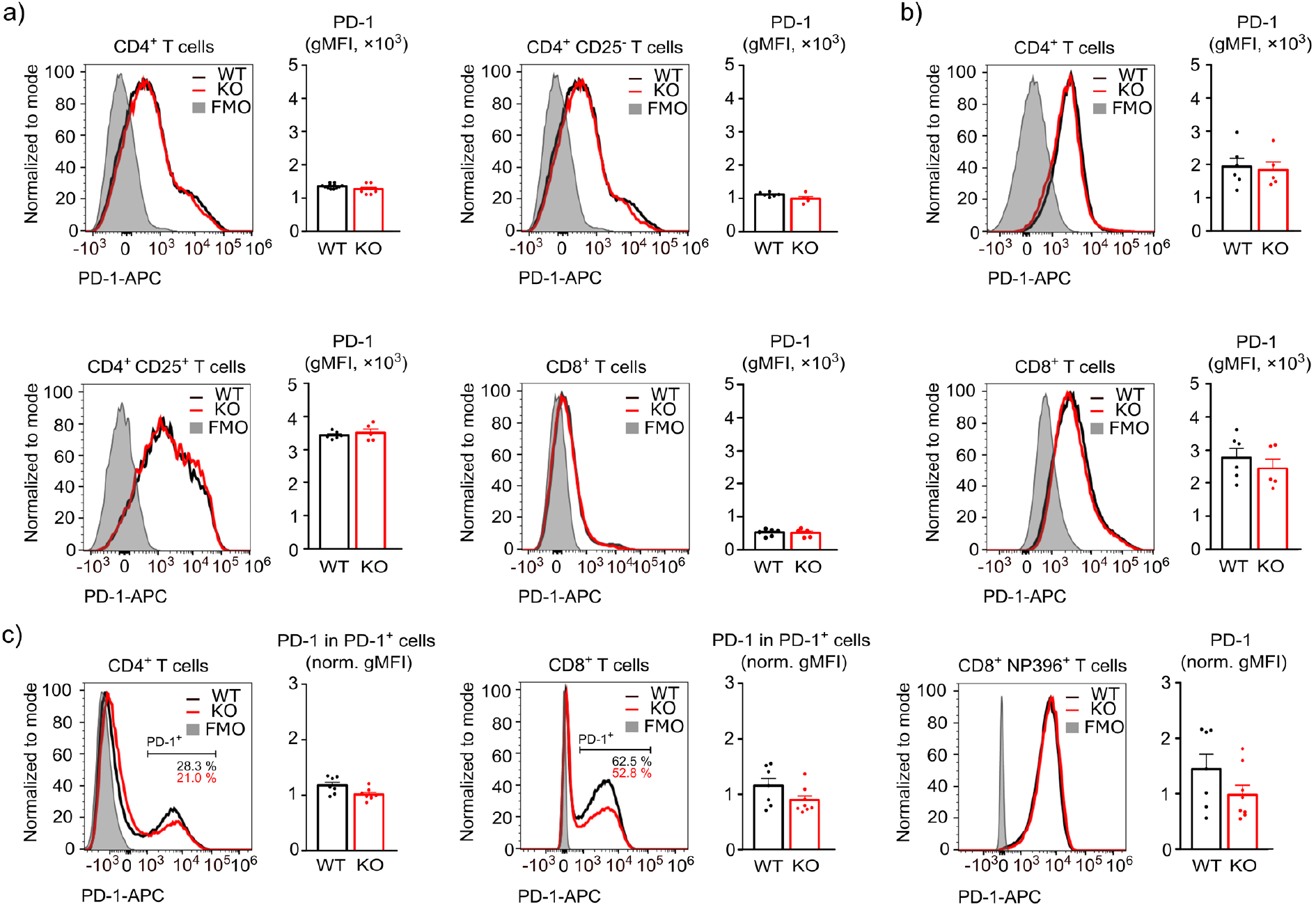
PD-1 surface levels are not altered in FBXO38-deficient T cells. (a,b) Cell-surface expressions of PD-1 in T cells isolated from the spleens of *Fbxo38*^*WT/WT*^ (WT) and *Fbxo38*^*KO/KO*^ (KO) mice. Histograms of representative mice and the quantification of the geometric mean fluorescence intensities (gMFI) for all mice (mean + SEM) are shown. (a) Steady-state surface levels of PD-1 in naïve CD4^+^, CD4^+^ CD25^-^, CD4^+^ CD25^+^, and CD8^+^ T cells. *n=* 6 *Fbxo38*^*WT/WT*^ and 5 *Fbxo38*^*KO/KO*^ mice in two independent experiments. (b) Surface levels of PD-1 in CD4^+^ (upper) and CD8^+^ T (lower) cells stimulated with anti- CD3/CD28 beads in the presence of IL-2 for 96 h. *n=* 6 *Fbxo38*^*WT/WT*^ and 5 *Fbxo38*^*KO/KO*^ mice in two independent experiments. (c) Cell-surface expressions of PD-1 in T cells isolated from the spleens of LCMV-infected *Fbxo38*^*WT/WT*^ and *Fbxo38*^*KO/KO*^ mice. The results of two independent experiments were normalized to the PD-1 gMFI of *Fbxo38*^*WT/KO*^ T cells (set as 1). Histograms represent results from littermate mice. Bars show normalized cell-surface PD-1 levels in CD4^+^ PD-1^+^, CD8^+^ PD-1^+^ and CD8^+^ D^b^-NP396 tetramer^+^ T cells (all PD-1^+^ gate). *n= 7 Fbxo38*^*WT/WT*^, 8 *Fbxo38*^*KO/KO*^, and 8 *Fbxo38*^*WT/KO*^ mice in two independent experiments. Individual experiments are shown in Extended Data Fig. 2.

*In vivo*, T-cell activation was induced by the lymphocytic choriomeningitis virus (LCMV) infection (Fig. 2c, Extended Data Fig. 2c-e). A relatively large fraction of CD4^+^ and CD8^+^ T cells upregulated PD-1 during the LCMV infection (Extended Data Fig. 2c). Again, PD-1 surface levels on PD-1^+^ CD4^+^ or CD8^+^ T cells were equal or slightly lower in *Fbxo38*^*KO/KO*^ than in WT mice (Fig. 2c). Using D^b^-NP396 tetramer, we gated on LCMV- specific CD8^+^ T cells. These cells were almost all PD-1 positive. Similar to the previous results, we observed comparable surface PD-1 expression in CD8^+^NP396^+^ tetramer^+^ T cells from WT and *Fbxo38*^*KO/KO*^ mice (Fig. 2c and Extended Data Fig. 2d,e).

Taken together, our data from both *in vivo* and cell systems do not support the conclusion that SCF^FBXO38^ is a ubiquitin ligase that regulates the level and the activity of PD-1 in T cells.

## Supporting information

Densitometric analysis of western blots shown in Figure 1 and in the Extended Data Figure 1.

Reagents used in the study.

## Extended Data Figure Legends

**Extended Data Fig. 1.**
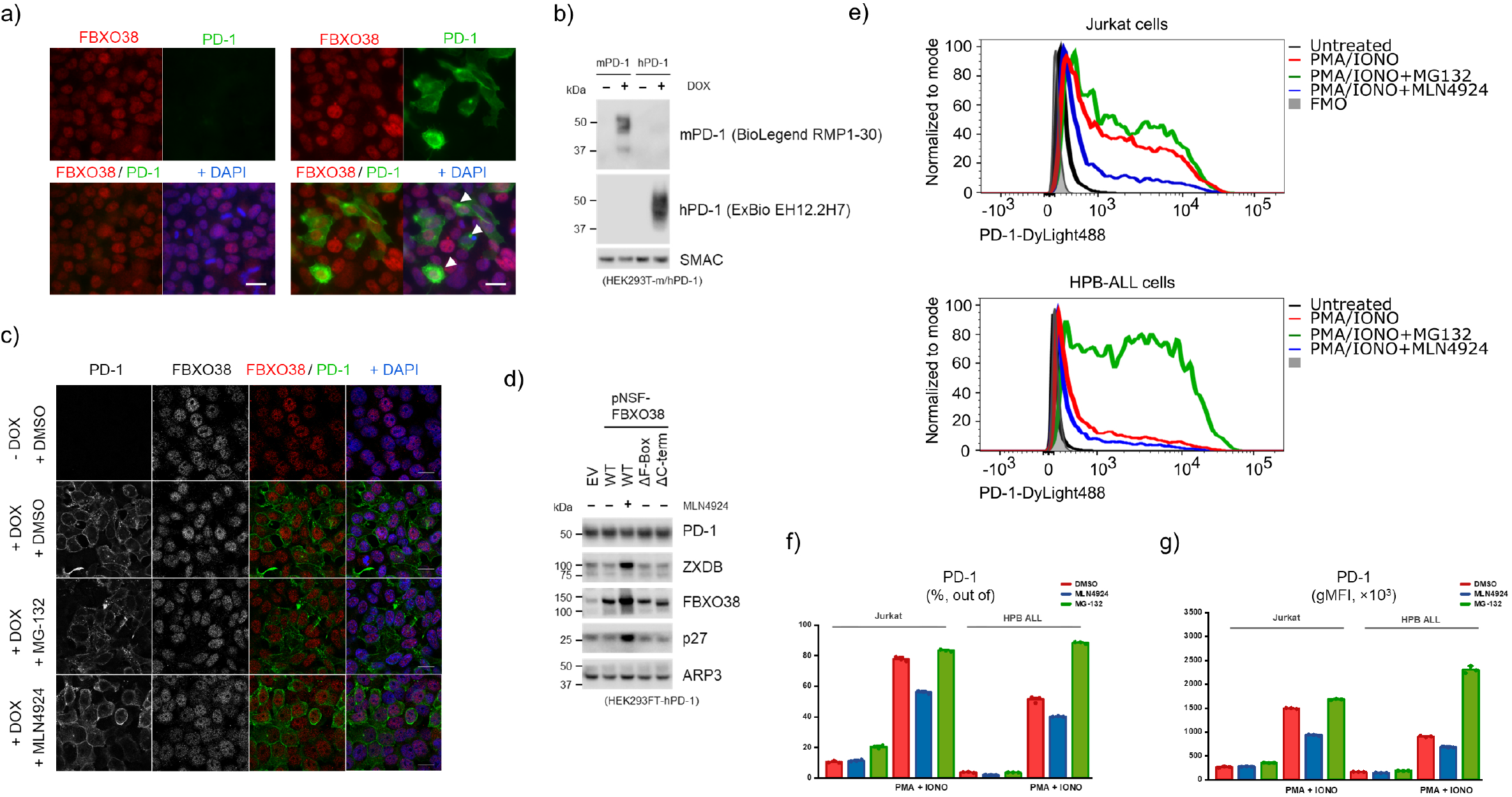
FBXO38 does not control PD-1 in the cell system. (a) Overexpressed PD-1 does not co-localize with endogenous FBXO38. HEK293FT cells were transfected with a pcDNA3.1 vector expressing human PD-1. The cells were fixed and permeabilized 48 h after transfection and subsequently incubated with FBXO38 and PD-1 antibodies. DNA was stained with DAPI. White arrowheads mark PD-1 cytosolic accumulation. Scale bar, 20 µm. (b) HEK293FT-mPD-1 and HEK293FT-hPD-1 cells were incubated with doxycycline, lysed, and immunoblotted as indicated. SMAC staining was used as a loading control. (c) HEK293FT-hPD-1 cells were incubated with doxycycline and then treated with MG-132 or MLN4924. Cells were fixed 6 h after treatments, permeabilized, and incubated with FBXO38 and PD-1 antibodies. DNA was stained with DAPI. Scale bar, 50 µm. (d) HEK293FT-hPD-1 cells were transfected with StrepII-FLAG-tagged wild-type FBXO38 (pNSF-FBXO38 WT), FBXO38 lacking the F-box motif (ΔF-Box) or C-terminus (ΔC-term). Where stated, the cells were treated with MLN4924 inhibitor 6 h prior to collection. Cells were lysed and immunoblotted as indicated. ZXDB or p27 staining was used as the positive control, and ARP3 as a loading control. (e-g) Jurkat or HPB-ALL T-cell leukemia cell lines were incubated for 72 h with phorbol myristyl acetate (PMA) and ionomycin (IONO) followed by 6 h treatments with MG-132 or MLN4924 (*n=* 3). Surface PD-1 expression was analyzed by flow cytometry. (e) Histograms represent the analysis of representative samples. (f,g) Graphs represent the percentage of PD-1-positive cells (f) and the geometric mean of PD-1 fluorescence intensities (gMFI + SD) from all cells (g).

**Extended Data Fig. 2.**
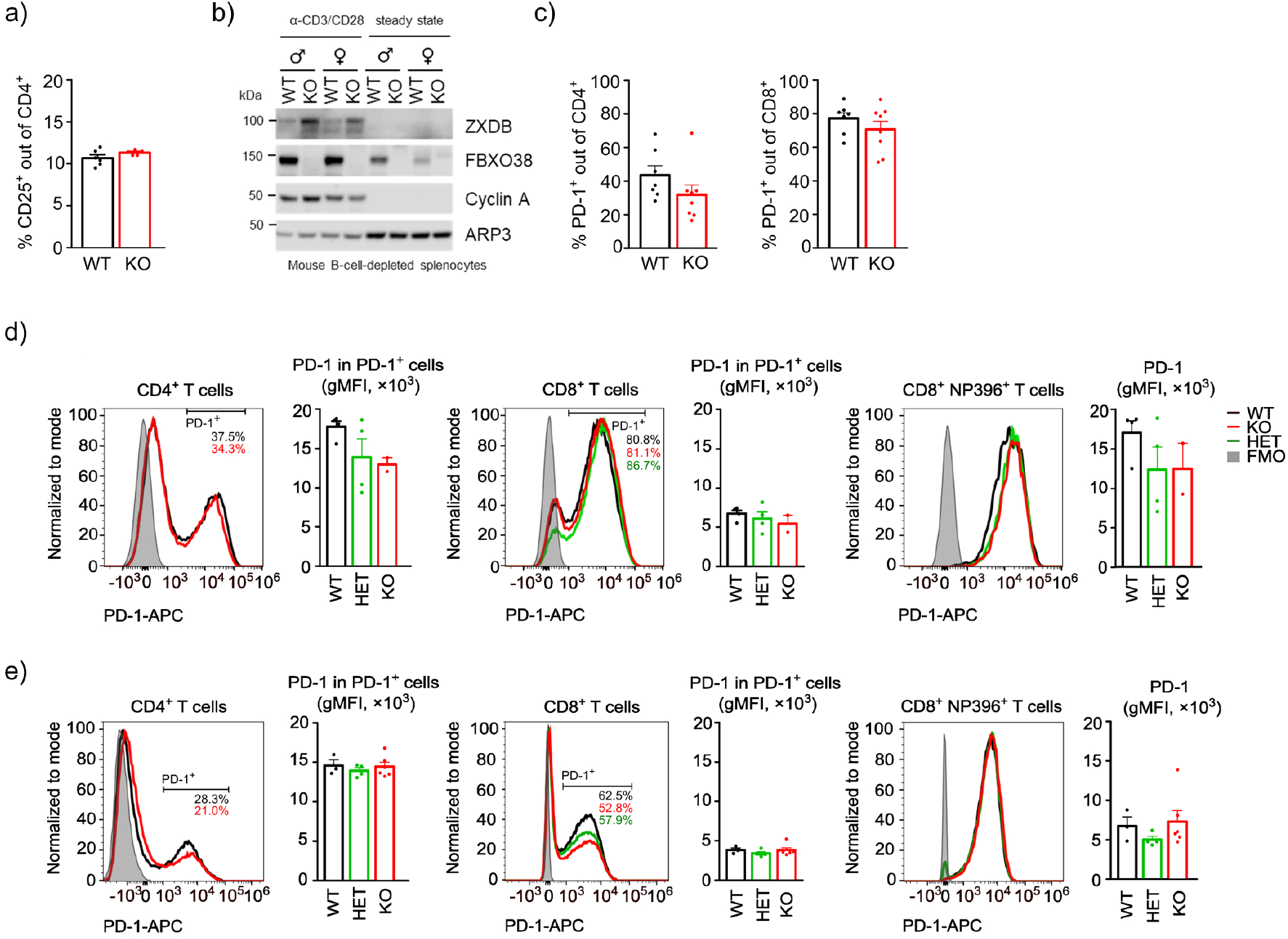
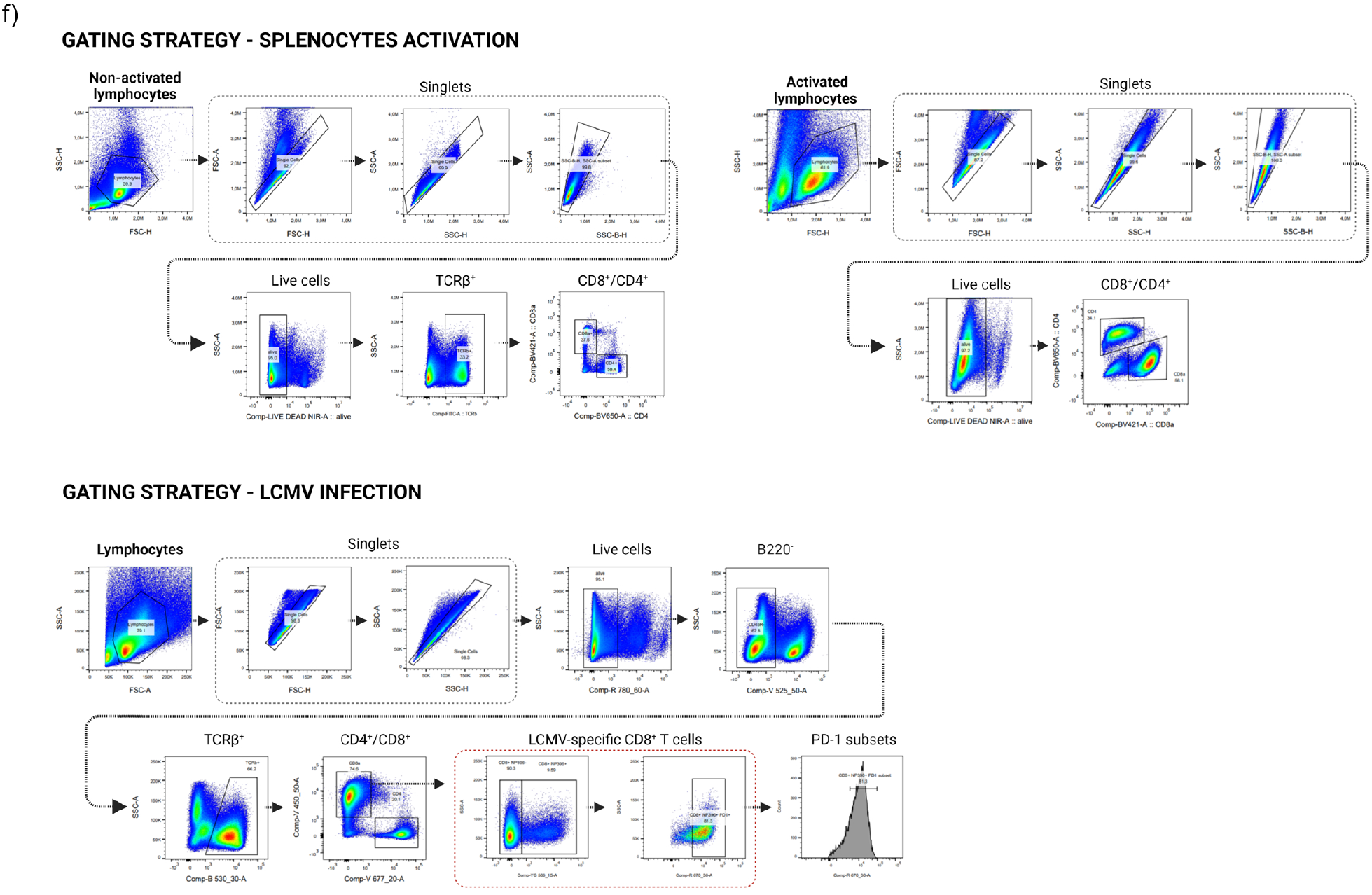
Analysis of FBXO38-deficient T cells. (a) Percentage of CD25^+^ cells out of CD4^+^ T cells. The bars represent mean + SEM. *n=* 6 *Fbxo38*^*WT/WT*^ (WT) and 5 *Fbxo38*^*KO/KO*^ (KO) mice in two independent experiments (same as in Fig. 2a). (b) Naïve or anti-CD3/CD28 stimulated T cells (same as in Fig. 2a, 2b) were lysed and immunoblotted as indicated. ZXDB or cyclin A staining was used as the positive control and ARP3 as a loading control. (c) Percentage of PD-1^+^ cells out of CD4^+^ and CD8^+^ T cells from *Fbxo38*^*WT/WT*^ and *Fbxo38*^*KO/KO*^ mice infected with LCMV. The bars represent mean + SEM. *n= 7 Fbxo38*^*WT/WT*^, 8 *Fbxo38*^*KO/KO*^ mice in two independent experiments (same as in Fig. 2c). (d,e) Cell-surface expressions of PD-1 in T cells isolated from the spleens of LCMV-infected *Fbxo38*^*WT/WT*^ and *Fbxo38*^*KO/KO*^ mice. The results of two independent experiments (d, e) are presented as normalized data in Fig. 2c. Histograms represent the analysis of representative samples. Bars show cell-surface PD-1 levels in CD4^+^ PD-1^+^, CD8^+^ PD-1^+^and CD8^+^ D^b^-NP396 tetramer^+^ T cells (all PD-1^+^ gate). *n= 7 Fbxo38*^*WT/WT*^, 8 *Fbxo38*^*KO/KO*^, and 8 *Fbxo38*^*WT/KO*^ mice. (f) Gating strategy used to analyze the flow cytometry data in Fig. 2 and Extended Data Fig. 2.

## Methods

### Cell culture procedures

Human cell line HEK293FT (ATCC #CRL-1573) was maintained in Dulbecco′s Modified Eagle′s Medium (DMEM) supplemented with 10% fetal bovine serum (FBS), and penicillin, streptomycin, and gentamicin, and cultured in a humidified incubator at 37 °C with 5% CO_2_. Where indicated, cycloheximide (100 µg/ml), MLN4924 (1µM; 6 h), MG-132 (10µM; 6 h), or doxycycline hyclate (0.2 µg/ml; 48 h) were used. Cells were transfected with pSBtet containing PD-1 and the transposase-containing pSB100X using Lipofectamine 2000 (Invitrogen) according to the manufacturer’s protocol. Positive clones were selected by puromycin 48 hours after transfection. Transient transfections were carried out using polyethylenimine (PEI MW 25000, Polysciences). Jurkat E6 (ATCC #TIB-152) and HPB-ALL (DSMZ #ACC-483) were cultured in RPMI 1640 medium supplemented with 10% FBS and antibiotics.

### Plasmid construction

cDNAs of human and mouse PDCD1 were amplified by PCR from human peripheral blood and mouse lymph node cDNAs, respectively. Prepared cDNAs were cloned into the pSBtet-Pur plasmid using SfiI restriction sites.

FBXO38-containing plasmids were created as described previously^3^. Briefly, human FBXO38 cDNA (Origene #RC204380) was cloned into pcDNA3.1 containing N-terminal twinStrepII- FLAG-tag (NSF). For the F-box-lacking FBXO38 variant, PCR mutagenesis was carried out using PfuX7 DNA polymerase with subsequent DpnI digestion.

### Gene silencing

Small interfering RNAs (Sigma Aldrich) were transfected into the subconfluent HEK293FT cells using Lipofectamine RNAiMAX (Invitrogen) according to the manufacturer’s protocol. Non-targeting siRNAs were used as negative controls.

### Immunoblotting

For whole-cell lysates, cells were washed with ice-cold PBS and lysed in a lysis buffer (150 mM NaCl, 50 mM Tris pH 7.5, 0.4% Triton X-100, 2 mM CaCl_2_, 2 mM MgCl_2_, 1 mM EDTA) in the presence of phosphatase a protease inhibitor cocktail (Pierce) and Benzonase nuclease (0.125 U/μl; Santa Cruz) for 30 min on ice. Lysates were mixed with an equal volume of 2% SDS in 50mM Tris-HCl, pH 8, heated for 5 min at 95 °C, and cleared by centrifugation. Protein concentration was determined by the BCA method (Thermo Fisher Scientific). Samples were prepared by mixing with Bolt™ LDS Sample Buffer (Thermo Fisher Scientific) supplemented with 10% β-mercaptoethanol (Sigma Aldrich), heated for 5 min at 95 °C, and then separated using NuPAGE™ 4-12% gradient Bis-Tris gel (Invitrogen). Afterward, the samples were transferred to PVDF membrane (Amersham), blocked with 5% non-fat milk (ChemCruz), and incubated with indicated antibodies diluted in 3% BSA (PanReac Applichem) in tris-buffered saline with Tween® 20 (TBST) overnight at 4 °C. The HRP-conjugated secondary antibodies (Cell Signaling) were diluted in 5% milk in TBS-T. The membranes were developed using WesternBright ECL (Advansta) or SuperSignal™ West Femto Maximum Sensitivity Substrate (Thermo Fisher Scientific).

### Protein affinity purification and immunoprecipitation

The cells were collected and lysed in the lysis buffer (150 mM NaCl, 50 mM Tris pH 7.5, 0.4% Triton X-100, 2 mM CaCl_2_, 2 mM MgCl_2_, 1 mM EDTA, supplemented with phosphatase and protease inhibitors) for 10 min on ice. For Strep-tagged FBXO38, the lysates cleared by centrifugation were incubated with Strep-Tactin® Sepharose resin (IBA Lifesciences). Purified proteins were then eluted by desthiobiotin using Buffer E (IBA Lifesciences) and subsequently prepared for immunoblotting as described above. For immunoprecipitation of PD-1 protein from 1×10^7^ HEK293FT-PD-1 cells, lysates were incubated with the PD-1 antibody (1 µg; Exbio #11-176) and mixed with Dynabeads® Protein G. Parental HEK293FT cells were used as a negative control. Elution of immunoprecipitated proteins was carried out with 1x Bolt™ LDS Sample Buffer (Thermo Fisher Scientific), and samples were prepared for immunoblotting as described above.

### Immunocytochemistry

Cells were washed with PBS, fixed with 3% PFA in PBS for 20 minutes, permeabilized with 0.2% Triton X-100 in PBS for 10 minutes, and blocked for 1 hour (3% BSA, 0.1% Triton X-100 in PBS). Incubation with indicated primary antibodies was carried out for 2 hours at RT, followed by incubation with Alexa Fluor-conjugated secondary antibodies (Abcam) for 30 minutes at RT. DAPI was used to stain DNA. Slides were mounted with ProLong Gold Antifade Mountant (Invitrogen). Images were acquired using Axio Imager Zeiss 2 (EC Plan-Nefluar objectives) and analyzed with ZEN 2.3 or ImageJ software.

### Mouse Model

*Fbxo38*^*KO/KO*^ mice were generated in a C57BL/6N background using the CRISPR/Cas9 genome-editing system as described^4^. Shortly, the Cas9 protein and gene-specific sgRNAs (Integrated DNA Technologies) were used for zygote electroporation. The genome editing was confirmed in the founder mouse using PCR.

### *In vitro* activation assay

T-cells were isolated from mouse splenocytes using erythrocyte lysis (ACK buffer), followed by B220^+^ or CD19^+^ B-cells depletion with specific antibodies and anti-rat magnetic beads (Invitrogen 11417D). 2×10^6^ of the remaining cells were stimulated with 2×10^6^ anti-CD3/CD28 beads (Gibco, 11453D) in 5 ml of IMDM medium supplemented with 10 % FBS, antibiotics, and 10 ng/ml IL-2 (Gibco) for 96 hours at 37 LCMV infection and 5 % CO_2_.

### LCMV infection

Mice were infected intraperitoneally with LCMV (Armstrong) dose of 2×10^5^ PFU. The spleens were harvested on day 8 post-infection. Splenocytes were depleted of erythrocytes by incubation in ACK buffer. The samples were analyzed using Cytek Aurora (Cytek) or FACSSymphony (BD Biosciences) flow cytometers.

### Flow cytometry analysis

For the analysis of mouse T cells, the following antibodies were used: anti-CD8α-BV421 (clone 53-6.7, BioLegend 100753, diluted 200×), anti-CD4-BV650 (clone RM4-5, BioLegend 100546, diluted 200×), anti-TCRβ-FITC (clone H57-597, BioLegend 109206, diluted 400×), anti-CD25-PE-Cy7 (clone PC61, BioLegend 102016, diluted 200×) or anti-CD25-PE (clone PC61, BioLegend 102008, diluted 200×), anti-PD-1-APC (clone 29F.1a12, BioLegend 135209, diluted 200×), anti-B220-BV510 (clone RA3-6B2, BioLegend 103248, diluted 200×). LIVE/DEAD fixable near-IR dye (Thermo Fisher Scientific, L34976) was used for the viability staining.

LCMV-specific CD8^+^ T cells were identified by specific H-2d^b^-NP396 tetramer staining (diluted 200×). The tetramers were produced by incubating biotinylated H-2D^b^-NP396-PE (FQPQNGQFI) monomers (from NIH Tetramer Core Facility) with streptavidin-PE (Thermo Fisher Scientific, S866) at a molar ratio of 3:1. Streptavidin-PE was added in two doses with 1 h incubation on ice after each step. Staining was performed in 2 % FBS, 2 mM EDTA, 0.1 % sodium azide in PBS on ice for 30 min. Full minus one (FMO) stained samples without anti-PD-1 antibody were used to gate PD-1^+^ cells.

For the analysis of human T cell-leukemia cell lines, anti-PD-1 (clone EH12-2H7; Exbio 11- 176, diluted 500×) in combination with anti-mouse IgG secondary (Thermo Fisher Scientific 35503; DyLight 488) was used. Propidium iodide (1 µg/ml in PBS; Sigma) was used for the viability staining.

### Quantification and Statistical Analysis

Graphs were generated using OriginPro 2021 and Prism 5.0 (GraphPad Software). The quantity/densitometry of protein in western blots was analyzed using Image Lab software (Bio-Rad). Flow cytometry data were analyzed using FlowJo version 10.6.2 (BD Biosciences). A representative experiment out of three is shown for all western blots and immunofluorescence pictures. The geometric mean was used for the bar charts, and the error bars show the standard deviation or standard error of the mean.

## Data Availability Statement

All original uncut blots, FACS data, and immunofluorescence files will be available in the Mendeley database after the publication of the manuscript.

## Ethics Statement

All animal experiments were approved by the Resort Professional Commission for Approval of Projects of Experiments on Animals of the Czech Academy of Sciences, Czech Republic (protocols 115/2016 and AVCR 1667/2022 SOV II) and were in accordance with the Czech Act No. 246/1992 Coll. and European directive 2010/63/EU.

Animals were maintained in a controlled, specific pathogen-free environment at the Animal Facilities of the Institute of Molecular Genetics of the Czech Academy of Sciences, Czech Republic. Mice were provided food and water *ad libitum* and kept in the animal facility with a 12-hour light-dark cycle.

## Author Contributions

Conceptualization, LC, ND, MP, OS; investigation, LC, ND, ES, KK; writing-original draft preparation, LC, ND; writing-review and editing, LC, ND, MP, OS; methodology, LC, ND, OS; supervision, LC, OS, MP; funding acquisition, LC, OS. All authors have read and agreed to the published version of the manuscript.

## Acknowledgments

We thank the Light Microscopy Core Facility, Institute of Molecular Genetics of the Czech Academy of Sciences, for their support with the imaging presented herein. In addition, we thank Dr. Tomas Brdicka for providing leukemia cell lines and antibody against SHP2. Illustrations were created with BioRender.com.

