## Supplementary material for "FBXO38 does not control PD-1 stability": Densitometric analysis of western blots shown in Figure 1 and in the Extended Data Figure 1.

| | | RAW DENSITOMETRY VALUES | | | | | NORMALIZED DENSITOMETRY VALUES (% OF CON.) | | | | |
|---|---|---|---|---|---|---|---|---|---|---|---|
| **Figure 1d** | **DOX.** | **PD-1** | **PARP** | **ARP3** | **FBXO38** | **p27** | **PD-1** | **PARP** | **ARP3** | **FBXO38** | **p27** |
| non-treated | + | 6101858 | 2638230 | 4824040 | 499310 | 724841 | 100 | 100 | 100 | 100 | 100 |
| MG-132 3 h | + | 6268304 | 2254322 | 4788511 | 845026 | 1470196 | 103 | 85 | 99 | 169 | 203 |
| MG-132 6 h | + | 5846568 | 2320514 | 4699162 | 893802 | 1819467 | 96 | 88 | 97 | 179 | 251 |
| MLN4924 3 h | + | 7021854 | 2418234 | 3129074 | 814926 | 1930838 | 115 | 92 | 65 | 163 | 266 |
| MLN4924 6 h | + | 6208916 | 2198434 | 2401503 | 756728 | 2199977 | 102 | 83 | 50 | 152 | 304 |
| non-treated | - | 205646 | 2041760 | 3440905 | 625030 | 472732 | 3 | 77 | 71 | 125 | 65 |

**Figure 1d (Ponceau S)**

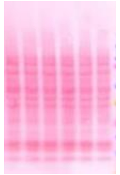

| | | RAW DENSITOMETRY VALUES | | | | | | NORMALIZED DENSITOMETRY VALUES (% OF CON.) | | | | | |
|---|---|---|---|---|---|---|---|---|---|---|---|---|---|
| **Figure 1e** | **DOX.** | **PD-1** | **PARP** | **ARP3** | **FBXO38** | **FBXO28** | **ZXDB** | **PD-1** | **PARP** | **ARP3** | **FBXO38** | **FBXO28** | **ZXDB** |
| control siRNA | + | 5879398 | 2332232 | 4847297 | 580734 | 725855 | 229600 | 100 | 100 | 100 | 100 | 100 | 100 |
| control siRNA 3 h CHX | + | 5022122 | 2904342 | 4329663 | 495656 | 422994 | 192388 | 85 | 125 | 89 | 85 | 58 | 84 |
| control siRNA 6 h CHX | + | 4491676 | 2439836 | 4210687 | 420980 | 536445 | 134582 | 76 | 105 | 87 | 72 | 74 | 59 |
| si FBXO38 siRNA | + | 5343548 | 2352182 | 4611685 | 83804 | 605111 | 945980 | 91 | 101 | 95 | 14 | 83 | 412 |
| si FBXO38 siRNA 3 h CHX | + | 4766034 | 2322152 | 4657809 | 30380 | 470470 | 667702 | 81 | 100 | 96 | 5 | 65 | 291 |
| si FBXO38 siRNA 6 h CHX | + | 3913280 | 2363298 | 5088980 | 42448 | 322504 | 549752 | 67 | 101 | 105 | 7 | 44 | 239 |
| si FBXO38 siRNA + NSF-FBXO38 | + | 5551294 | 1798678 | 4749251 | 1726018 | 746694 | 629181 | 94 | 77 | 98 | 297 | 103 | 274 |
| si FBXO38 siRNA + NSF-FBXO38 3 h CHX | + | 5437964 | 1989442 | 4730206 | 1624364 | 568087 | 375053 | 92 | 85 | 98 | 280 | 78 | 163 |
| si FBXO38 siRNA + NSF-FBXO38 6 h CHX | + | 3862026 | 2215458 | 4014348 | 1220744 | 353665 | 309309 | 66 | 95 | 83 | 210 | 49 | 135 |

**Figure 1e (Ponceau S)**

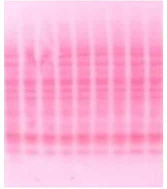

| | | RAW DENSITOMETRY VALUES | | | | | NORMALIZED DENSITOMETRY VALUES (% OF CON.) | | | | |
|---|---|---|---|---|---|---|---|---|---|---|---|
| **Extended Data Figure 1d** | **DOX.** | **PD-1** | **ARP3** | **FBXO38** | **ZXDB** | **p27** | **PD-1** | **ARP3** | **FBXO38** | **ZXDB** | **p27** |
| non-transfected | + | 6159384 | 5026424 | 533764 | 1035160 | 908791 | 100 | 100 | 100 | 100 | 100 |
| NSF-FBXO38 WT | + | 6350848 | 4468828 | 1921514 | 777952 | 993902 | 103 | 89 | 360 | 75 | 109 |
| FNSF-BXO38 WT + MLN4924 | + | 5639186 | 4995081 | 3094154 | 1647954 | 1846026 | 92 | 99 | 580 | 159 | 203 |
| NSF-FBXO38 ΔF-Box | + | 6617226 | 4660305 | 1982498 | 866796 | 1016626 | 107 | 93 | 371 | 84 | 112 |
| NSF-FBXO38 ΔC-term | + | 6502804 | 4810572 | 2401854 | 812882 | 847288 | 106 | 96 | 450 | 79 | 93 |

**Extended Data Figure 1d (Ponceau S)**

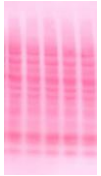
