## Supplementary material for "FBXO38 does not control PD-1 stability": Reagents used in the study.

| Name | Company | CAT# | Concentration |
|---|---|---|---|
| 2-mercaptoethanol | Sigma | M6250 | |
| 4',6-diamidin-2-fenylindol (DAPI) | Sigma | D9542 | |
| Benzonase | Santa Cruz | sc-391121 | 0.125 U/µl |
| Cycloheximide | Sigma | C7698 | 100 µg/ml |
| Desthiobiotin | IBA Lifesciences | 2-1000-025 | 2.5 mM |
| Dithiothreitol | Sigma | 10197777001 | |
| Doxycycline hyclate | Sigma | D9891 | 0.2 µg/ml |
| Formaldehyde | Thermo Fisher Scientific | 28908 | |
| Gentamycin | Sandoz | | 40 mg/ml |
| Il2 | Gibco | PMC0024 | 10 ng/ml |
| Ionomycin calcium salt | Sigma | 10634 | 1 µM |
| Lipofectamine 2000 | Thermo Fisher Scientific | 11668019 | |
| LIVE/DEAD fixable near-IR-dye | Thermo Fisher Scientific | L34976 | |
| Methanol | Penta | 67-56-1 | |
| MG132 | Medchemexpresss | HY-13259 | 10 µM |
| MLN4924 | Medchemexpresss | HY-70062 | 1 µM |
| Paraformaldehyde | Electron Microscopy Sciences | 15710 | |
| Penicilin | BB Pharma | | 100 U/ml |
| Phorbol myristyl acetate (PMA) | Sigma | P1585 | 30 ng/ml |
| Polyethyleneimine (MW 25 000) | Polysciences | 23966 | |
| Ponceau S | VWR | K793 | |
| Propidium iodide | Sigma | 537059 | 1 µg/ml |
| Protease Inhibitors Mini Tablets | Pierce | A32955 | |
| Proteinase K | Sigma | P2308 | |
| Puromycin | Sigma | P8833 | 1 µg/ml |
| RNAiMax | Thermo Fisher Scientific | 13778150 | |
| Sodium dodecyl sulfate (SDS) | Sigma | 71736 | |
| Sodium fluoride | Sigma | S7920 | |
| Sodium orthovanadate | Sigma | 450243 | |
| Streptomycin | Sigma | S9137 | 100 mg/ml |
| Triton X-100 | Sigma | T8787 | |

**Cloning primers**

| Gene | Orientation | Position | Sequence | RE SITE |
|---|---|---|---|---|
| hPD-1 | FWD | FL | AAAGGCCTCTGAGGCCACCATGCAGATCCCACAGGCGCCCTGG | SFI1 |
| hPD-1 | REV | FL | CTTGGCCTGACAGGCCTCAGAGGGGCCAAGAGCAGTGTCC | SFI1 |
| mPD-1 | FWD | FL | AAAGGCCTCTGAGGCCACCATGTGGGTCCGGCAGGTACCCTGG | SFI1 |
| mPD-1 | REV | FL | CTTGGCCTGACAGGCCTCAAAGAGGCCAAGAACAATGTCC | SFI1 |
| hFBXO38 | FWD | FL | CGAGAAAGGAGCTAGCGGGCCACGAAAGAAAAGTG | NHE1 |
| hFBXO38 | REV | Fl | AATCCTCTCCGCTAGCTTAAATGTAGTCATCTTCAACT | NHE1 |
| hFBXO38_ΔC-term | REV | 1000 | GGGTCGACAAGTCTTCTTCACTTCGAGT | SAL1 |

**Mutagenesis primers**

| Gene | Mutation | Position | Sequence |
|---|---|---|---|
| FBXO38 | (ΔFBOX) | Δ30-65 | TCTCACAACTCGCAGATATAGATAGTCCTTTGTTTCATCTGC |

**CRISPR**

| FBXO38 CRISPR targeting sequences | Orientation | Position | Sequence |
|---|---|---|---|
| FBXO38_5' | Upstream | Exon 3 | TTAAGTAAATACAGGTAAGCCGG |
| FBXO38_3' | Downstream | Exon 3 | GGGTGTGCTGTTTAGAGGGCAGG. |
| **CRISPR PCR verifiction primers** | **Orientation** | **Position** | **Sequence** |
| mFBXO38 | FWD | Exon 3 | GGCTGTTCTCCTCTTTGTG |
| mFBXO38 | REV | Exon 3 | TCCTTCCTCCTGCTAGACTCT |

| Primary antibodies | Source | CAT# | RRID | LINK | Clone | Dilution |
|---|---|---|---|---|---|---|
| ARP3 | Santa Cruz | ab49671 | AB_2257830 | AB_2257830 | | |
| SH-PTP2 | Santa Cruz | sc-280 | AB_632401 | AB_632401 | | |
| FBXO28 | Bethyl | A302-377A | AB_1907260 | AB_1907260 | | |
| FBXO38 | Atlas Antibodies | HPA041444 | AB_2677484 | AB_2677484 | | |
| FLAG | Cell Signaling | 14793 | AB_2572291 | AB_2572291 | | |
| p27 | Santa Cruz | sc-528 | AB_632129 | AB_632129 | | |
| PARP-1 | Santa Cruz | sc-8007 | AB_628105 | AB_628105 | | |
| SKP1 | Cell Signaling | 12248 | AB_2754993 | AB_2754993 | | |
| Strep II Tag | Novus Biologicals | NBP2-43735 | AB_2916323 | AB_2916323 | | |
| ZXDA/B | Atlas Antibodies | HPA043789 | AB_2678673 | AB_2678673 | | |
| mPD-1 | Bio Legend | 109111 | AB_10613470 | AB_10613470 | RMP1-30 | |
| hPD-1 | ExBio | 11-176 | AB_2687629 | AB_2687629 | EH12.2H7 | |
| Cyclin A | Homemade (Dr. M. Pagano) | | | | | |
| B220 | BioLegend | 103204 | AB_312989 | AB_312989 | RA3-6B2 | 25x |
| CD19 | BioLegend | 152402 | AB_2629714 | AB_2629714 | 1D3 | 25x |

| Fluorescent antibodies / proteins | Source | CAT# | RRID | LINK | Clone | Dilution |
|---|---|---|---|---|---|---|
| CD8α-BV421 | BioLegend | 100753 | AB_2562558 | AB_2562558 | 53-6.7 | 200x |
| anti-CD4-BV650 | BioLegend | 100546 | AB_2562098 | AB_2562098 | RM4-5 | 200x |
| anti-TCRβ-FITC | BioLegend | 109206 | AB_313429 | AB_313429 | H57-597 | 400x |
| anti-CD25-PE | BioLegend | 102008 | AB_312857 | AB_312857 | PC61 | 200x |
| anti-CD25-PE-Cy7 | BioLegend | 102016 | AB_312865 | AB_312865 | PC61 | 200x |
| anti-PD-1-APC | BioLegend | 135209 | AB_2251944 | AB_2251944 | 29F.1A12 | 200x |
| Anti-B220-BV510 | BioLegend | 103248 | AB_2650679 | AB_2650679 | RA3-6B2 | 200x |
| Streptavidin, R-Phycoerythrin Conjugate (SAPE) | Thermo Fisher Scientific | S866 | | | | |

| Secondary antibodies | Source | CAT# | RRID | LINK |
|---|---|---|---|---|
| Anti-Rabbit IgG (Alexa Fluor® 555) | Abcam | ab150070 | AB_2783636 | AB_2783636 |
| Anti-Human IgG (Alexa Fluor 488) | Thermo Fisher Scientific | A11013 | AB_2534080 | AB_2534080 |
| anti-Mouse IgG (DyLight 488) | Thermo Fisher Scientific | 35503 | AB_1965946 | AB_1965946 |
| Anti-mouse IgG, HRP-linked | Cell signaling | 7076 | AB_330924 | AB_330924 |
| Anti-rabbit IgG, HRP-linked | Cell signaling | 7074 | AB_2099233 | AB_2099233 |

| Beads for protein purification | Source | CAT# |
|---|---|---|
| Strep-Tactin® Superflow resin | IBA | 2-1206-025 |
| Protein G (magnetic) | Dynabeads | 10004D |
| Dynabeads Untouched Mouse CD8 Cells Kit | Thermo Fisher Scientific | 11417D |

| Beads for activation | Source | CAT# |
|---|---|---|
| Dynabeads® Mouse T-Activator CD3/CD28 | Thermo Fisher Scientific | 11453D |

| Tertamer | Source |
|---|---|
| Biotinylated MHC I monomer H-2Db-NP396 (FQPQNGQFI) | NIH Tetramer Core Facility |

| Expression vectors | Backbone | N-term. tag |
|---|---|---|
| FBXO38 | pCDNA3 (pNSF) | 1xFLAG-2xStrepII |
| hPD-1 | pCDNA3 | |
| hPD-1 | pSB | |
| mPD-1 | pSB | |

| Sleeping Beauty System | Company | Cat. Number |
|---|---|---|
| pSBtet-Pur | Addgene | 60507 |
| pSB100X | Addgene | 34879 |

| Name | Sequence | Target | Position | CAT# |
|---|---|---|---|---|
| FBXO38#1 | GGGUGUAUUUCAGCGAGUAUU | FBXO38 | TR | |
| FBXO38#2 | GGACUCGAUUGGUUGAUAUUU | FBXO38 | TR | |
| FBXO38#3 | GAGCGAAGCUGUUUGAGUAUU | FBXO38 | UTR | |
| CTRL_#1 | AUGAACGUGAAUUGCUCAAUU | NON-TARG | | D-001210-04 |
| CTRL_#2 | AUGUAUUGGCCUGUAUUAGUU | NON-TARG | | D-001210-03 |
